# Estimating bobcat (*Lynx rufus*) abundance in Wisconsin during 1981-2015

**DOI:** 10.1101/2020.03.23.004044

**Authors:** Maximilian L. Allen, Nathan Roberts, Javan Bauder

## Abstract

Quantifying and estimating trends in wildlife abundance is critical for their management and conservation. Harvest-based indices are often used as a surrogate index for wildlife population. Sex-Age-Kill (SAK) models generally use the age-at-harvest of males and females, combined with annual mortality and reproduction rates to calculate a preharvest population estimate. We used and SAK model to estimate abundance for bobcats from 1981-2015. Pre-hunt population size ranged from approximately 1630-2148 during 1981-1995 after which the population increased to a maximum of 4439 in 2005 before declining to 2598 in 2013. Pre-hunt population size was highly correlated an index of abundance from winter track counts (*r* = 0.93). We found that the model, as currently implemented by WNDR, appears to provide an accurate trend of statewide bobcat abundance. SAK models more logistically feasible for long-term evaluations of population trends overbroad spatial extents than more intensive methods. While SAK models may be the only technique available to wildlife managers for estimating the abundance of harvested species, we encourage additional research to evaluate the effects of potential biases on estimates of abundance.

## Introduction

Estimating trends in wildlife abundance is critical for their management, but this can be difficult for cryptic species (Gese 2001, Allen et al. 2018a). Hence, indices, including harvest-based indices, are often used as a surrogate for wildlife populations (Gese 2001, Skalski et al. 2005). Harvest records may span multiple decades, and often provide the only long-term data source for populations in a given management unit (McCullough et al. 1990, Newsome and Ripple 2015). There are many models currently available that estimate abundance using harvest data, and the model used often depends on the type of data available.

One population model used by agencies, including the Wisconsin Department of Natural Resources (WDNR) are sex-age-kill (SAK) models (Creed et al. 1984, Roseberry and Woolf 1991). SAK models are retroactive and dependent on accurate harvest data (Mattson and Moritz 2008). SAK models generally combines the age-at-harvest of males and females, combined with annual mortality and reproduction rates to calculate a preharvest population estimate (Creed et al. 1984). These population estimates can be for entire states or smaller management units. Population size in subsequent years is then calculated based on the previous year’s demographics and current year’s age- and sex-specific harvest rates.

The WDNR has a long history of using SAK for white-tailed deer (*Odocoileus virginianus*) management (Creed et al. 1984), but also uses SAK to help manage other species including American black bear (*Ursus americanus*) (Allen et al. 2018a) and bobcat (*Lynx rufus*) (Allen et al. 2019). We used and SAK model to estimate abundance for bobcats from 1981– 2015.

## Methods

The WDNR has a mandatory harvest registration for bobcats. As part of registration, the skinned carcasses of all harvested bobcats must be submitted to the WDNR. We aged every harvested bobcat annually using the cementum annuli on an extracted canine tooth (Crowe 1972). We counted the of placental scars and uteri counts from all harvested female bobcats each year to estimate annual pregnancy rates. for yearling and adult females. We assumed constant mean annual litter sizes of 2.5 for yearlings and 2.7 for adults based on the mean counts of placental scars and uteri.

We assumed a constant sex ratio of 46% males and initial age distributions of 36% juveniles, 23% yearlings, and 41% adults (Erb 2009). We assumed conservative estimates of non-harvest mortality rates for yearlings (summer = 0.35, winter = 0.30, and annual = 0.545) and adult males and females (summer = 0.13, winter = 0.13, and annual 0.24) (Roberts and Dennison 2017). We also estimated age- and sex-specific harvest rates from data on the age and sex of registered harvested bobcats. We assumed that mortality from poaching (i.e., unregistered harvest) was 0.10 from 1981–1988 and 0.20 thereafter when harvest became more restricted (Allen et al. 2018b).

We evaluated the correlation between estimated abundance and an independent index of abundance estimated from winter track counts conducted by WDNR since 1977 in 20 counties in northern Wisconsin (Crimmins and Van Deelen 2019). Two 16-km tracks were surveyed in each county with 8–37 visits per track. Tracks were placed along remote roads with minimal human use and surveyed during November and December. See Crimmins and Van Deelen (Crimmins and Van Deelen 2019) for additional details on survey methodology. We used the mean maximum number of bobcat tracks seen per route.

### Statistical Model

We calculated initial pre-birth population size in each age/sex class by multiplying the initial population size, the overall proportion of that sex, and the proportion of that age class (Fig. 1). We calculated initial post-birth population size as the current year’s number of females in each age class multiplied by the age-specific estimate of pregnancy rate and litter size and 0.5 (assuming equal sex ratio at birth). Post-birth population size for yearlings was the pre-birth population size of juveniles, while the post-birth population size for adults was the sum of pre-birth population size of yearlings and adults.

**Figure 1.**
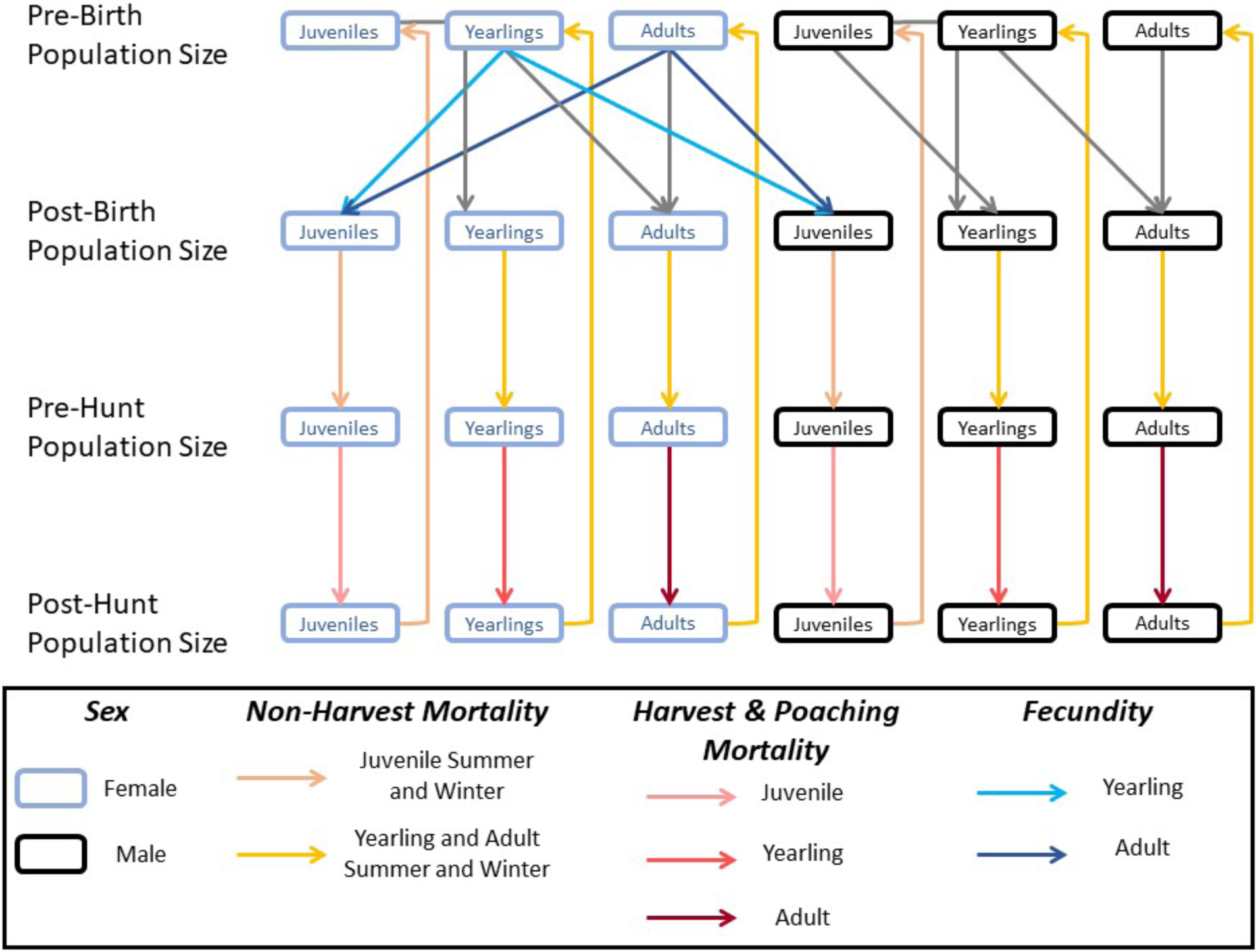
Flow diagram of the sex-age-kill model used to estimate bobcat abundance in Wisconsin. Fecundity includes age-specific estimate of pregnancy rate and litter size and an equal sex ratio at birth between males and females.

We calculated summer non-harvest mortality for each age/sex class as post-birth population size and summer non-harvest mortality rate. We then calculated pre-hunt population size as post-birth population size minus summer non-harvest mortality. We calculated harvest and poaching mortality for each age/sex class as the sum of the registered harvest for each sex plus the estimated number killed by poaching (1 + poaching rate) times the proportion of harvest for that class. We calculated winter non-harvest mortality as described for summer non-harvest mortality.

We calculated post-harvest population size as pre-harvest population size minus harvest and poaching mortality. The subsequent year’s pre-birth population size for each age/sex group was calculated as post-harvest population size minus winter non-harvest mortality. Our abundance estimate was the sum of pre-hunt population size across all age/sex groups. We started the model at 1981 and ran the model through the 2015–2016 season using an initial population size of 1,430 based on historical estimates (Rolley et al. 2001, Roberts and Dennison 2017).

## Results

Mean pregnancy rates were 0.34 for yearlings and 0.69 for adults (Table 1). A mean of 98 female and 141 male bobcats were harvested annually (Table 2). The proportions of females harvested by age class were 0.22, 0.23, and 0.55 for juveniles, yearlings, and adults, respectively. The proportions of males harvested by age class were 0.22, 0.21, and 0.57 for juveniles, yearlings, and adults, respectively.

**Table 1.** Reproductive parameters estimated from harvested bobcats (*Lynx rufus*) in Wisconsin during 1981–2015 for use in a sex-age-kill model to estimate population size.

| Year | Yearling | Yearling | 2-Year Old | 2-Year Old | Adult | Adult |
| --- | --- | --- | --- | --- | --- | --- |
|  | Pregnancy Rate | Litter Size | Pregnancy Rate | Litter Size | Pregnancy Rate | Litter Size |
| 1981 | 0.380 | 2.5 | 0.620 | 2.7 | 0.620 | 2.7 |
| 1982 | 0.380 | 2.5 | 0.620 | 2.7 | 0.620 | 2.7 |
| 1983 | 0.250 | 2.5 | 0.773 | 2.7 | 0.773 | 2.7 |
| 1984 | 0.111 | 2.5 | 0.588 | 2.7 | 0.588 | 2.7 |
| 1985 | 0.313 | 2.5 | 0.556 | 2.7 | 0.556 | 2.7 |
| 1986 | 0.333 | 2.5 | 0.704 | 2.7 | 0.704 | 2.7 |
| 1987 | 0.400 | 2.5 | 0.790 | 2.7 | 0.790 | 2.7 |
| 1988 | 0.722 | 2.5 | 0.938 | 2.7 | 0.938 | 2.7 |
| 1989 | 0.571 | 2.5 | 0.696 | 2.7 | 0.696 | 2.7 |
| 1990 | 0.273 | 2.5 | 0.700 | 2.7 | 0.700 | 2.7 |
| 1991 | 0.200 | 2.5 | 0.667 | 2.7 | 0.667 | 2.7 |
| 1992 | 0.381 | 2.5 | 0.706 | 2.7 | 0.706 | 2.7 |
| 1993 | 0.143 | 2.5 | 0.400 | 2.7 | 0.400 | 2.7 |
| 1994 | 0.273 | 2.5 | 0.750 | 2.7 | 0.750 | 2.7 |
| 1995 | 0.625 | 2.5 | 0.688 | 2.7 | 0.688 | 2.7 |
| 1996 | 0.714 | 2.5 | 0.923 | 2.7 | 0.923 | 2.7 |
| 1997 | 0.417 | 2.5 | 0.872 | 2.7 | 0.872 | 2.7 |
| 1998 | 0.294 | 2.5 | 0.750 | 2.7 | 0.750 | 2.7 |
| 1999 | 0.571 | 2.5 | 0.906 | 2.7 | 0.906 | 2.7 |
| 2000 | 0.438 | 2.5 | 0.717 | 2.7 | 0.717 | 2.7 |
| 2001 | 0.625 | 2.5 | 0.692 | 2.7 | 0.692 | 2.7 |
| 2002 | 0.539 | 2.5 | 0.902 | 2.7 | 0.902 | 2.7 |
| 2003 | 0.410 | 2.5 | 0.860 | 2.7 | 0.860 | 2.7 |
| 2004 | 0.318 | 2.5 | 0.740 | 2.7 | 0.740 | 2.7 |
| 2005 | 0.079 | 2.5 | 0.817 | 2.7 | 0.817 | 2.7 |
| 2006 | 0.320 | 2.5 | 0.667 | 2.7 | 0.667 | 2.7 |
| 2007 | 0.083 | 2.5 | 0.667 | 2.7 | 0.667 | 2.7 |
| 2008 | 0.000 | 2.5 | 0.410 | 2.7 | 0.410 | 2.7 |
| 2009 | 0.401<sup>1</sup> | 2.5 | 0.563 | 2.7 | 0.563 | 2.7 |
| 2010 | 0.000 | 2.5 | 0.542 | 2.7 | 0.542 | 2.7 |
| 2011 | 0.090 | 2.5 | 0.710 | 2.7 | 0.710 | 2.7 |
| 2012 | 0.143 | 2.5 | 0.489 | 2.7 | 0.489 | 2.7 |
| 2013 | 0.159<sup>2</sup> | 2.5 | 0.647 | 2.7 | 0.647 | 2.7 |
| 2014 | 0.159<sup>2</sup> | 2.5 | 0.561 | 2.7 | 0.561 | 2.7 |
| 2015 | 0.159<sup>2</sup> | 2.5 | 0.584<sup>3</sup> | 2.7 | 0.584<sup>3</sup> | 2.7 |

<sup>1</sup>Value was not recorded during that year so the mean value during 1997–2006 was used.
<sup>2</sup>Values were not recorded during those years so the mean value during 2004–2012 was used.
<sup>3</sup>Values were recorded during that year so the mean values during 2006–2014 were used.

**Table 2.** *Harvest parameters for bobcat (*Lynx rufus*) in Wisconsin during 1981–2015 for use in a sex-age-kill model to estimate population size.*

| Year | Total Reported Harvest |  | Proportion of Juveniles |  | Proportion of Yearlings |  | Proportion of Adults |  | Poaching |
| --- | --- | --- | --- | --- | --- | --- | --- | --- | --- |
|  | Females | Males | Females | Males | Females | Males | Females | Males | Rate |
| 1981 | 96 | 112 | 0.280 | 0.280 | 0.310 | 0.280 | 0.410 | 0.440 | 0.10 |
| 1982 | 64 | 75 | 0.280 | 0.280 | 0.310 | 0.280 | 0.410 | 0.440 | 0.10 |
| 1983 | 107 | 99 | 0.133 | 0.359 | 0.378 | 0.231 | 0.489 | 0.410 | 0.10 |
| 1984 | 121 | 139 | 0.295 | 0.226 | 0.227 | 0.283 | 0.477 | 0.491 | 0.10 |
| 1985 | 91 | 98 | 0.286 | 0.357 | 0.393 | 0.214 | 0.321 | 0.429 | 0.10 |
| 1986 | 78 | 105 | 0.290 | 0.270 | 0.304 | 0.258 | 0.406 | 0.472 | 0.10 |
| 1987 | 107 | 140 | 0.340 | 0.302 | 0.223 | 0.294 | 0.436 | 0.405 | 0.10 |
| 1988 | 80 | 85 | 0.417 | 0.284 | 0.300 | 0.313 | 0.283 | 0.403 | 0.10 |
| 1989 | 65 | 71 | 0.235 | 0.231 | 0.275 | 0.250 | 0.490 | 0.519 | 0.20 |
| 1990 | 50 | 48 | 0.500 | 0.375 | 0.283 | 0.175 | 0.217 | 0.450 | 0.20 |
| 1991 | 32 | 39 | 0.308 | 0.226 | 0.231 | 0.323 | 0.462 | 0.452 | 0.20 |
| 1992 | 86 | 131 | 0.227 | 0.208 | 0.307 | 0.264 | 0.467 | 0.528 | 0.20 |
| 1993 | 73 | 87 | 0.222 | 0.265 | 0.270 | 0.191 | 0.508 | 0.544 | 0.20 |
| 1994 | 57 | 112 | 0.269 | 0.218 | 0.192 | 0.267 | 0.538 | 0.515 | 0.20 |
| 1995 | 44 | 67 | 0.219 | 0.213 | 0.250 | 0.149 | 0.531 | 0.638 | 0.20 |
| 1996 | 68 | 98 | 0.256 | 0.274 | 0.186 | 0.258 | 0.558 | 0.468 | 0.20 |
| 1997 | 105 | 111 | 0.194 | 0.278 | 0.194 | 0.167 | 0.612 | 0.556 | 0.20 |
| 1998 | 83 | 111 | 0.324 | 0.191 | 0.243 | 0.223 | 0.432 | 0.585 | 0.20 |
| 1999 | 77 | 111 | 0.321 | 0.315 | 0.125 | 0.157 | 0.554 | 0.528 | 0.20 |
| 2000 | 108 | 171 | 0.165 | 0.240 | 0.200 | 0.163 | 0.635 | 0.597 | 0.20 |
| 2001 | 60 | 92 | 0.186 | 0.134 | 0.186 | 0.164 | 0.628 | 0.701 | 0.20 |
| 2002 | 120 | 130 | 0.167 | 0.119 | 0.167 | 0.191 | 0.667 | 0.691 | 0.20 |
| 2003 | 143 | 228 | 0.111 | 0.215 | 0.172 | 0.101 | 0.717 | 0.684 | 0.20 |
| 2004 | 154 | 208 | 0.198 | 0.232 | 0.191 | 0.196 | 0.611 | 0.572 | 0.20 |
| 2005 | 184 | 313 | 0.160 | 0.214 | 0.271 | 0.140 | 0.570 | 0.646 | 0.20 |
| 2006 | 137 | 219 | 0.096 | 0.148 | 0.260 | 0.179 | 0.644 | 0.673 | 0.20 |
| 2007 | 178 | 299 | 0.244 | 0.196 | 0.153 | 0.210 | 0.603 | 0.593 | 0.20 |
| 2008 | 124 | 243 | 0.153 | 0.162 | 0.245 | 0.203 | 0.602 | 0.635 | 0.20 |
| 2009 | 89 | 182 | 0.141 | 0.170 | 0.155 | 0.163 | 0.704 | 0.667 | 0.20 |
| 2010 | 127 | 222 | 0.105 | 0.105 | 0.092 | 0.070 | 0.803 | 0.825 | 0.20 |
| 2011 | 146 | 211 | 0.023 | 0.073 | 0.148 | 0.163 | 0.830 | 0.764 | 0.20 |
| 2012 | 96 | 146 | 0.182 | 0.146 | 0.106 | 0.068 | 0.712 | 0.786 | 0.20 |
| 2013 | 86 | 140 | 0.133 | 0.081 | 0.222 | 0.135 | 0.645 | 0.784 | 0.20 |
| 2014 | 86 | 178 | 0.136 | 0.146 | 0.170 | 0.130 | 0.695 | 0.724 | 0.20 |
| 2015 | 83 | 169 | 0.116<sup>1</sup> | 0.110<sup>1</sup> | 0.148<sup>1</sup> | 0.113<sup>1</sup> | 0.737<sup>1</sup> | 0.777<sup>1</sup> | 0.20 |

<sup>1</sup>Values not recorded during those years so the mean values during 2010–2014 were used.

Pre-hunt population size ranged from approximately 1630–2148 during 1981–1995 after which the population increased to a maximum of 4439 in 2005 before declining to 2598 in 2013 (Figure 2). Pre-hunt population size was highly correlated the index of abundance from winter track counts (*r* = 0.78) and the 3-year running average of winter track counts (*r* = 0.93).

**Figure 2.**
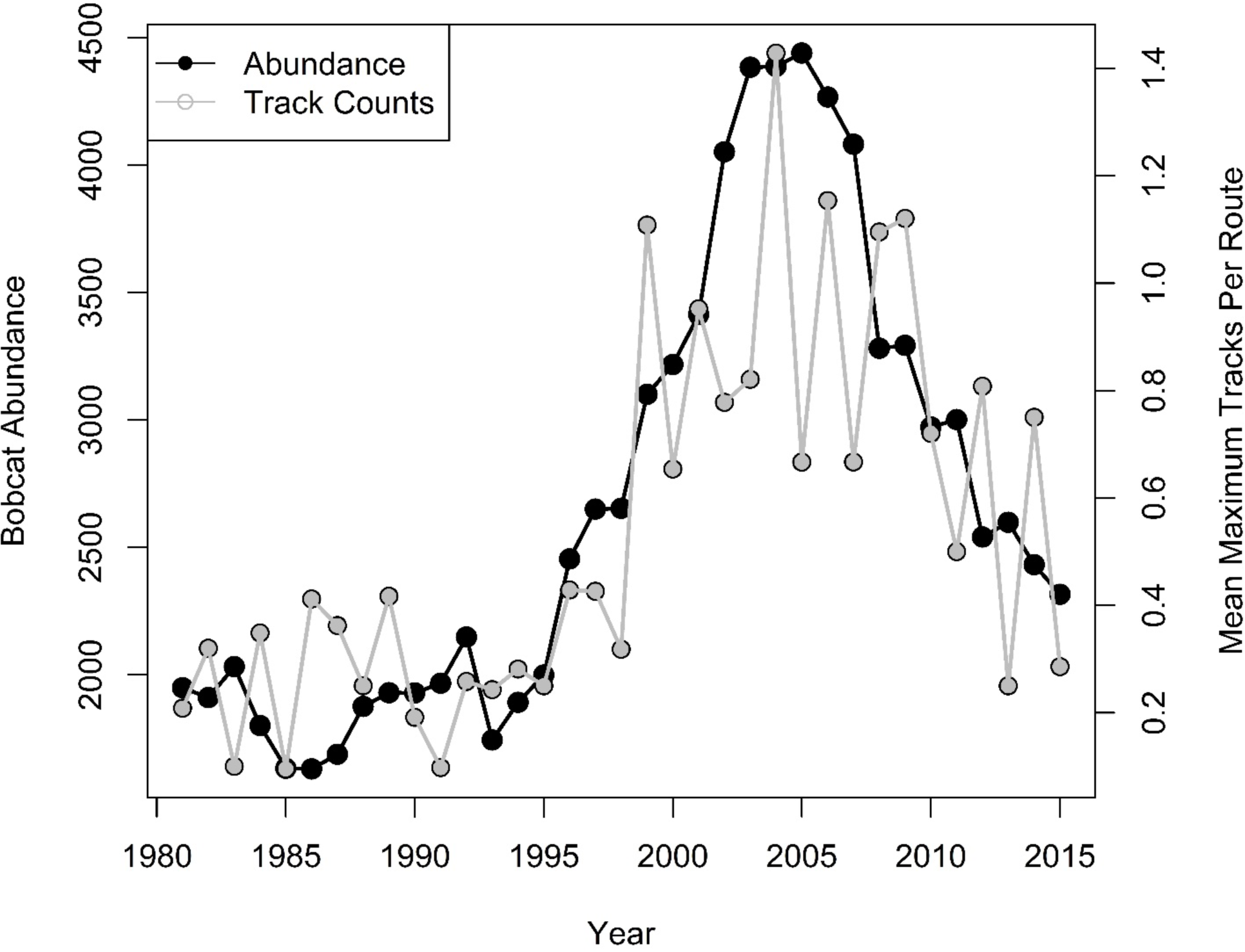
Estimated bobcat (Lynx rufus) population size in Wisconsin during 1981–2015 using a sex-age-kill model. The mean maximum number of bobcat tracks seen per route is shown for comparison.

## Discussion and Conclusions

The SAK we describe has been an integral part of WDNR’s bobcat management for many years (Roberts and Dennison 2017). We found that the model, as currently implemented by WNDR, appears to provide an accurate trend of statewide bobcat abundance. The data required for this model can be obtained annually through the requirement that successful bobcat hunters and trappers register their kill. This makes SAK models more logistically feasible for long-term evaluations of population trends overbroad spatial extents than more intensive methods such as radio telemetry or mark-recapture. Nevertheless, there are limitations to estimating abundance from SAK models including assumptions of stable age distributions and population growth (Millspaugh et al. 2009). However, Mattson and Moritz (Mattson and Moritz 2008) found that combining archery and firearm harvest of while-tailed deer did not alter population trends despite violating the assumption of stable age distributions. While SAK models may be the only technique available to wildlife managers for estimating the abundance of harvested species, we encourage additional research to evaluate the effects of potential biases on estimates of abundance.

